# Fast and Robust Inference of Phylogenetic Ornstein-Uhlenbeck Models Using Parallel Likelihood Calculation

**DOI:** 10.1101/115089

**Authors:** Venelin Mitov, Tanja Stadler

## Abstract

Phylogenetic comparative methods have been used to model trait evolution, to test selection versus neutral hypotheses, to estimate optimal trait-values, and to quantify the rate of adaptation towards these optima. Several authors have proposed algorithms calculating the likelihood for trait evolution models, such as the Ornstein-Uhlenbeck (OU) process, in time proportional to the number of tips in the tree. Combined with gradient-based optimization, these algorithms enable maximum likelihood (ML) inference within seconds, even for trees exceeding 10,000 tips. Despite its useful statistical properties, ML has been criticised for being a point estimator prone to getting stuck in local optima. As an elegant alternative, Bayesian inference explores the entire information in the data and compares it to prior knowledge but, usually, runs in much longer time, even for small trees. Here, we propose an approach to use the full potential of ML and Bayesian inference, while keeping the runtime within minutes. Our approach combines (i) a new algorithm for parallel likelihood calculation; (ii) a previously published method for adaptive Metropolis sampling. In principle, the strategy of (i) and (ii) can be applied to any likelihood calculation on a tree which proceeds in a pruning-like fashion leading to enormous speed improvements. As a showcase, we implement the phylogenetic Ornstein-Uhlenbeck mixed model (POUMM) in the form of an easy-to-use and highly configurable R-package. In addition to the above-mentioned usage of comparative methods, the POUMM allows to estimate non-heritable variance and phylogenetic heritability. Using simulations and empirical data from 487 mammal species, we show that the POUMM is far more reliable in terms of unbiased estimates and false positive rate for stabilizing selection, compared to its alternative - the non-mixed Ornstein-Uhlenbeck model, which assumes a fully heritable and perfectly measurable trait. Further, our analysis reveals that the phylogenetic mixed model (PMM), which assumes neutral evolution (Brownian motion) can be a very unstable estimator of phylogenetic heritability, even if the Brownian motion assumption is only weakly violated. Our results prove the need for a simultaneous account for selection and non-heritable variance in phylogenetic evolutionary models and challenge stabilizing selection hypotheses stated in numerous macro-evolutionary studies.

## Introduction

The past decades have seen active developement of phylogenetic comparative models of trait evolution, progressing from null neutral models, such as single-trait Brownian motion (BM), to complex multi-trait models incorporating selection, interaction between trait values and diversification, and co-evolution of multiple traits (O’Meara 2012; Manceau, Lambert, and Morlon 2016). Recent works have shown that, for a broad family of phylogenetic comparative models, the likelihood of an observed tree and data conditioned on the model parameters can be computed in time proportional to the size of the tree (FitzJohn 2012; Ho and Ańe 2014; Goolsby, Bruggeman, and Ańe 2016; Manceau, Lambert, and Morlon 2016). This family includes Gaussian models like Brownian motion and Ornstein-Uhlenbeck phylogenetic models as well as some non-Gaussian models like phylogenetic logistic regression (Emmanuel Paradis and Claude 2002; Ives and Garland 2009; Ho and Ańe 2014). All of these likelihood calculation techniques rely on post-order tree traversal or ‘pruning’ as coined by (Felsenstein 1973). Using pruning algorithms for likelihood calculation in combination with a gradient-based optimization method (Boyd and Vandenberghe 2004), maximum likelihood model inference runs within seconds on contemporary computers, even for phylogenies containing many thousands of tips (Ho and Ańe 2014). Other important features of the maximum likelihood estimate (MLE) are its simple interpretation as the point in parameter space maximizing the probability of the observed data under the assumed model, and its theoretical properties making it ideal for hypothesis testing and for model selection via likelihood ratio tests and information criteria. However, a major disadvantage of MLE is that, being a point estimate, it does not allow to explore the likelihood surface. Further, gradient based optimization, while fast, is prone to getting stuck in local optima.

As an elegant alternative, Bayesian approaches such as Markov Chain Monte Carlo (MCMC) provide posterior samples and high posterior density (HPD) intervals for the model parameters but require many orders of magnitude more likelihood evaluations. This represents a bottleneck in Bayesian analysis, in particular, when faced with large phylogenies of many thousands of tips, such as transmission trees from large-scale epidemiological studies, e.g. Hodcroft et al. (2014). While big data provides sufficient statistical power to fit a complex model, the time needed to perform a full scale Bayesian inference often limits the choice to a faster but less informative ML-inference, or a Bayesian inference on a simplified model. Another issue with Bayesian methods is that they require some level of expertise for specifying an apropriate prior, assessing the convergence of the MCMC and interpreting the results.

In this article, we propose a general approach allowing to use ML and Bayesian inference to their full potential, even for complex phylogenetic comparative models and for very large trees exceeding millions of tips, where the limiting factor becomes the available memory and not the calculation time. To achieve this goal, our approach combines two ideas: (i) the pruning algorithm for likelihood calculation can be accelerated by orders of magnitude through parallelization on modern multi-core processors and graphics adapters; (ii) the number of iterations needed for MCMC convergence can be reduced by the use of adaptive Metropolis sampling (Vihola 2012; Scheidegger 2012). Our parallel algorithm relies on a previously unexplored representation of the likelihood function as a quadratic polynomial of the trait value at the root. A nice property of the algorithm is that its parallel efficiency converges to 1 as the number of tips in the tree goes to infinity. Thus, for large trees, the parallel speed-up is practically limited by the number of available processing cores.

Numerous studies have discussed the Ornstein-Uhlenbeck (OU) process (Ornstein and Zernike 1919; Uhlenbeck and Ornstein 1930) as a model for trait adaptation under stabilizing selection, see e.g. Hansen (1997), Beaulieu et al. (2012), L. J. Harmon et al. (2010), Manceau, Lambert, and Morlon (2016) and references therein. However, in a cautionary note, Cooper et al. (2015) questionned the application of OU as a validation model for stabilizing selection and showed through simulations that OU-inferences are prone to overestimating the strength of selection if they do not account for measurement error. Connecting to these studies and providing a showcase for our parallel pruning algorithm, we implemented the phylogenetic Ornstein-Uhlenbeck mixed model (POUMM). Formally, POUMM can be regarded as adding a white noise term to the non-mixed phylogenetic OU model (POU), also referred to as single stationary peak (SSP) model (L. J. Harmon et al. 2010) and “Hansen” model with a single selection regime (Hansen 1997; Butler and King 2004). This white noise term is interpreted as a non-heritable component, i.e. a contribution to the measured trait-value not explainable by the assumed phylogenetic model, such as a measurement error, an environmental contribution and a model residual. Another way to interpret the POUMM is as an extension of the phylogenetic mixed model (PMM) (Lynch 1991; Housworth, Martins, and Lynch 2004), replacing the BM process with an OU process. The POUMM combines the applications of the above two models and, as we will show, is more reliable in terms of correct estimation of selection strength and phylogenetic heritability. However, currently, there are no software tools supporting fast Bayesian POUMM inference on large non-ultrametric trees. We provide our implementation in the form of a package written in the R language of statistical computing (R Core Team 2013). Based on our simulations, the time for a combined MLE and MCMC-fit on a tree of 10,000 tips, including two parallel MCMC chains of a million iterations is reduced from days to a few minutes. We present the model and its applications, the parallel algorithm for likelihood calculation and simulation results validating the technical correctness of the software and comparing its performance and robustness to alternative models and implementations. The POUMM R-package has already been used in several studies quantifying the heritability of continous traits, such as the “set-point virus load” and the “CD4 cell decline” in large HIV phylogenies with tips sampled through time (Mitov and Stadler (2016), Blanquart et al. (2017), Bertels et al. (2017), Bachmann et al. (2017)). Here, we additionally apply the method to an ultrametric tree and body weight data from 487 extant mammal species, including monophyletic groups of 227 Rodentia, 138 Chiroptera and 122 Soricomorpha species (Bininda-Emonds et al. 2007; Raia, Carotenuto, and Meiri 2010; Smith et al. 2003). Strikingly, the analysis of this data reveals that the outcome of model selection based on a likelihood ratio (LR) test and the Akaike information criterion (AICc) depends on the inclusion of a non-heritable component in the model. When comparing models with a non-heritable component (PMM vs POUMM), the Brownian-motion based PMM gets selected for the three mammal orders, as well as the combined phylogeny. Conversely, if comparing models assuming full heritablity, the POU model gets selected over non-mixed phylogenetic Brownian motion (PBM). For all trees the PMM model has the best AICc compared to all other models. These results challenge previous hypotheses of stabilizing selection that have been validated through POU models acting at the *class* and *order* levels (e.g. Raia and Meiri (2011)). Before us, others have pointed out this issue in simulation studies (Cooper et al. 2015). But the continuous use of phylogenetic comparative models assuming full phylogenetic heritability shows the low awareness for that issue and the need to provide strong empirical evidence. We revisit this issue in the Discussion section.

## Materials and Methods

Through the rest of the article we will rely on the following setup. Given is a rooted phylogenetic tree *𝒯* with *N* tips indexed by 1*, …, N* and a root node, 0 (Fig. 1). Without restrictions on the tree topology, non-ultrametric trees (i.e. tips have different time-distance from the root) and polytomies (i.e. nodes with any finite number of descendants) are accepted. Internal nodes are indexed by the numbers *N* + 1, …. Associated with the tips is a *N*-vector of observed real trait values denoted by **z**. We denote by *𝒯_i_* the subtree rooted at node *i* and by **z***_i_* the set of values at the tips belonging to *𝒯_i_*. For any internal node *j*, we denote by *Desc*(*j*) the set of its direct descendants. Furthermore, for any *i* ∈ *Desc*(*j*), we denote by *t_ji_* the length of the edge connecting *j* with *i* and by *t*_0_*_i_* the sum of edge-lengths (time-distance) from the root to *i*. The mean root-tip distance in the tree is denoted by 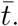 For two tips *i* and *k*, we denote by *t*_0(*ik*)_ the time-distance from the root to their most recent common ancestor (mrca), and by *τ_i_k* the sum of edge-lengths on the path from *i* to *k* (also called phylogenetic/patristic distance between *i* and *k*).

**Figure 1:**
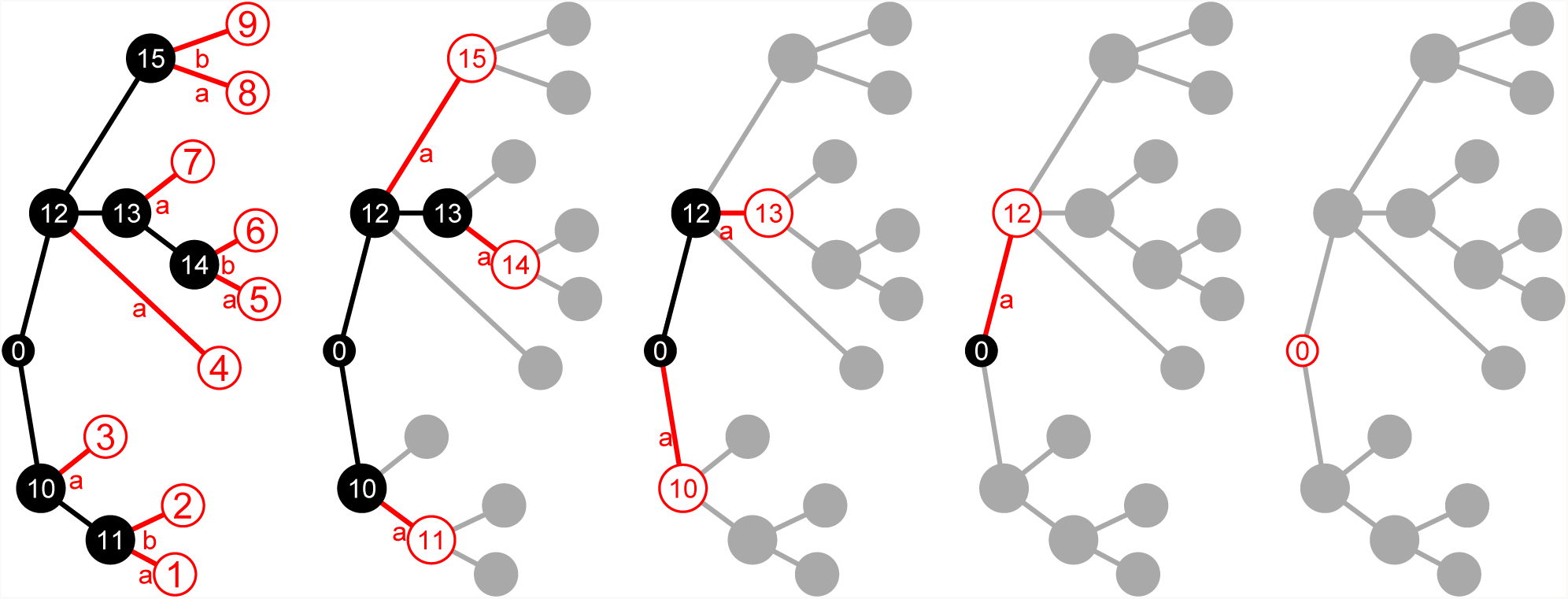
Breadth-first pruning on a tree with *N* = 9 tips. Each tree from left to right depicts one pruning iteration; black: non-tip nodes at a current pruning step; red: tip nodes to be pruned; grey: pruned nodes. Letters ‘a’ and ‘b’ next to branches denote the order in which the coefficients *a_ji_*, *b_ji_*, *c_ji_* are added to their parent’s *a_j_*, *b_j_* and *c_j_* (algorithm 1).

### The Phylogenetic Ornstein-Uhlenbeck Mixed Model

The phylogenetic Ornstein-Uhlenbeck mixed model (POUMM) decomposes the trait value as a sum of a non-heritable component, *e*, and a genetic component, *g*, which (i) evolves continuously according to an Ornstein-Uhlenbeck (OU) process along branches; (ii) gets inherited by the branches descending from each internal node. In biological terms, *g* is a genotypic value (Lynch and Walsh 1998) that evolves according to random drift with stabilizing selection towards a global optimum; *e* is a non-heritable component, which can be interpreted in different ways, depending on the application, i.e. a measurement error, an environmental contribution, a residual with respect to a model prediction, or the sum of all these. The OU-process acting on *g* is parameterized by an initial genotypic value at the root, *g*_0_, a global optimum, *θ*, a selection strength, *α>*0, and a random drift unit-time standard deviation, *σ*. Denoting by *W_t_* the standard Wiener process (Grimmett and Stirzaker 2001), the evolution of the trait-value, *z*(*t*), along a given lineage of the tree is described by the equations:

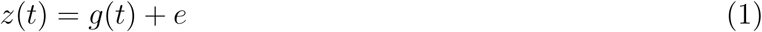

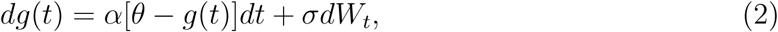

The stochastic differential equation 2 defines the OU-process, which represents a random walk tending towards the global optimum *θ* with stronger attraction for bigger difference between *g*(*t*) and *θ* (Ornstein and Zernike 1919; Uhlenbeck and Ornstein 1930). The model assumptions for *e* are that they are iid normal with mean 0 and standard deviation *σ_e_* at the tips. Any process along the tree that gives rise to this distribution at the tips may be assumed for *e*. For example, in the case of epidemics, a newly infected individual is assigned a new *e*-value which represents the contribution from its immune system and this value can change or remain constant throughout the course of infection. In the case of macro-evolution, *e* may represent the ecological (non-genetic) differences between species. In particular, the non-heritable component *e* does not influence the behavior of the OU-process *g*(*t*). Thus, if we were to simulate trait values *z* on the tips of a phylogenetic tree *𝒯*, we could first simlate the OU-process from the root to the tips to obtain *g*, and then add the white noise *e* (i.e. an iid draw from a normal distribution) to each simulated *g* value at the tips.

The POUMM represents an extension of the phylogenetic mixed model (PMM) (Lynch 1991; Housworth, Martins, and Lynch 2004), since, in the limit *α* → 0, the OU-process converges to a Brownian motion (BM) with unit-time standard deviation *σ*. Both, the POUMM and the PMM, define an expected multivariate normal distribution for the trait values at the tips. The mean vectors and the variance-covariance matrices of these distributions are written in table 1. Note that the trait expectation and variance for a tip *i* depends on its time-distance from the root (*t*_0_*_i_*), and the trait covariance for a pair of tips (*ij*) depends on the time-distance from the root to their mrca (*t*0(*ij*)), and, in the case of POUMM, on their patristic distance (*τ_ij_*) (table 1).

**Table 1:** Population properties at the tips of the phylogeny under POUMM and PMM. *µ_i_*: expected value at tip *i*; Σ*_ii_* : expected variance for tip *i*; Σ*_ij_* : expected covariance of the values of tips *i* and *j*; 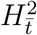: phylogenetic heritability at mean root-tip distance; 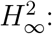: phylogenetic heritability at long-term equilibrium; 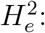: time-independent (empirical) phylogenetic heritability. Since the expressions for the expected variance-covariance matrix of the POUMM are only defined for strictly positive *α*, the expressions for PMM are obtained noting that 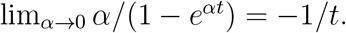.

| | POUMM | PMM ($\alpha \rightarrow 0$) |
| --- | --- | --- |
| $\Theta$: | $\langle g_0, \alpha, \theta, \sigma, \sigma_e \rangle$ | $\langle g_0, \sigma, \sigma_e \rangle$ |
| $\mu_i(\Theta, \mathcal{T})$: | $e^{-\alpha t_{0i}} g_0 + (1 - e^{-\alpha t_{0i}}) \theta$ | $g_0$ |
| $\Sigma_{ii}(\Theta, \mathcal{T})$: | $\frac{\sigma^2}{2\alpha} (1 - e^{-2\alpha t_{0i}}) + \sigma_e^2$ | $\sigma^2 t_{0i} + \sigma_e^2$ |
| $\Sigma_{ij}(\Theta, \mathcal{T})$: | $\frac{\sigma^2}{2\alpha} e^{-\alpha \tau_{ij}} (1 - e^{-2\alpha t_{0(ij)}})$ | $\sigma^2 t_{0(ij)}$ |
| $H_t^2$: | $\frac{\sigma^2(1 - e^{-2\alpha \bar{t}})}{\sigma^2(1 - e^{-2\alpha \bar{t}}) + 2\alpha\sigma_e^2}$ | $\bar{t}\sigma^2/(\bar{t}\sigma^2 + \sigma_e^2)$ |
| $H_\infty^2$: | $\sigma^2/(\sigma^2 + 2\alpha\sigma_e^2)$ | 1 |
| $H_e^2$: | $1 - \sigma_e^2/s^2(\mathbf{z})$ | $1 - \sigma_e^2/s^2(\mathbf{z})$ |

### A parallel pruning algorithm for fast likekelihood calculation

For a fixed tree, *𝒯*, the log-likelihood of the observed data is defined as the function:

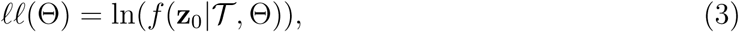

where *f* denotes a probability density function (pdf) and Θ =*< g*_0_*, α, θ, σ, σ_e_ >*. Here, we propose a parallel variant of the pruning algorithm (Felsenstein 1973). The log-likelihood is calculated by consecutive integration over the unobservable genotypic values, *g_i_*, progressing from the tips to the root. Central for the algorithm is the following theorem:

#### Theorem 1

(Quadratic polynomial representation of the POUMM log-likelihood). *For α* ≥ 0, *a real θ and non-negative σ and σ_e_, the POUMM log-likelihood can be expressed as a quadratic polynomial of g*_0_:

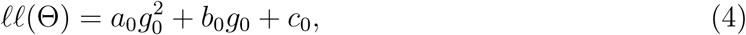

*where a*_0_ < 0, *b*_0_ *and c*_0_ *are real coefficients. We denote by u*(*α, t*) *the function:*

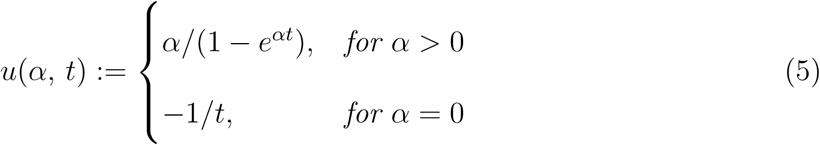

*Then, the coefficients in eq. 4 can be expressed with the following recurrence relation:*

1. *For j* ∈ {1, …, *N*} *(tips):*

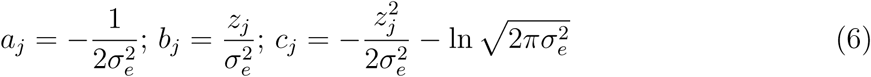
2. *For j* > *N (internal nodes) or j* = 0 *(root):*

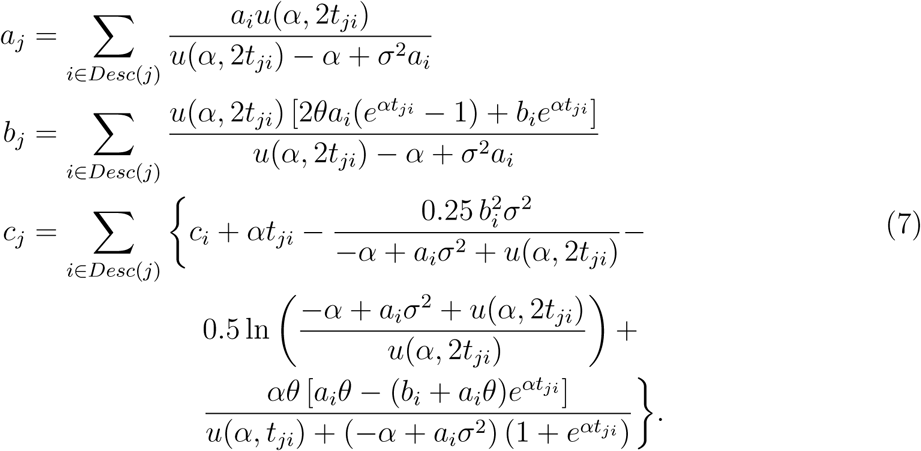

*Proof.* Induction from the tips to the root of the tree.

- *Basis:* For a tip-node *i*, *𝒯_i_* is the trivial tree consisting of this tip-node only and the pdf of **z***_i_*, conditioned on the unobservable genotypic value *g_i_*, is given by the normal pdf with mean *g_i_* and standard deviation *σ_e_*. This pdf can be written as:

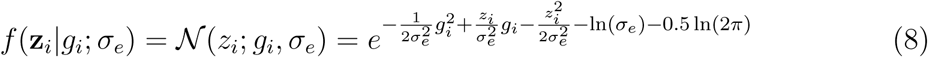 By defining 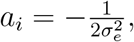, 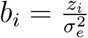 and 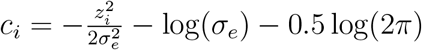 and taking the natural logarithm of the pdf we obtain the log-likelihood representation from eq. 4, where *a*_0_ < 0, *b*_0_ and *c*_0_ can be calculated from the observed value *z_i_* and the model parameter *σ_e_*.
- *Inductive hypothesis:* Assume that for an internal node *j*, the statement of the theorem has been proven for all subtrees *𝒯_i_*, *i* ∈ *Desc*(*j*).
- *Inductive step:* Substituting *g_j_* for *g*_0_ and *t_ji_* for *t*_0_*_i_* in the POUMM expressions for *µ_i_* and Σ*_ii_* (Table 1), and integrating over *g_i_*, we can write the pdf of **z***_i_*, conditioned on *g_j_*, *t_ji_* and Θ:

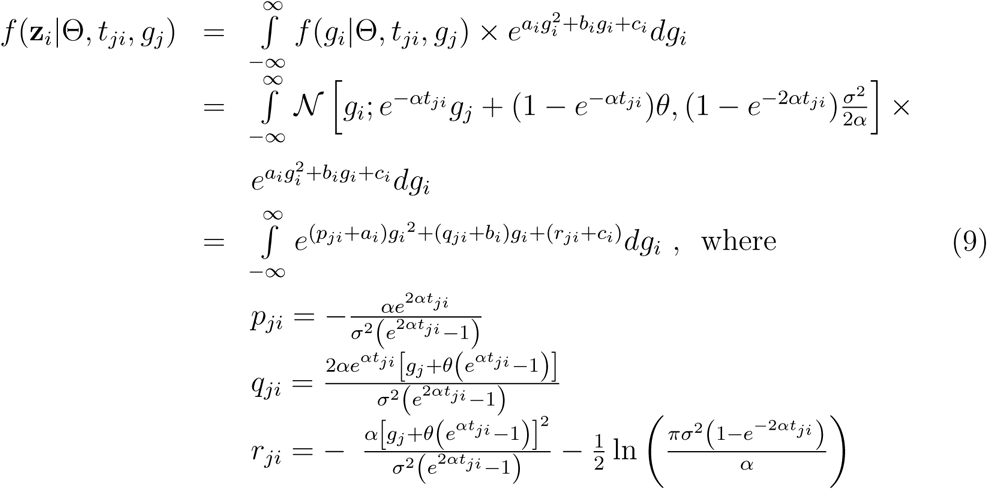 We notice that *p_ji_*, *q_ji_* and *r_ji_* in eq. 9 are not defined in the case of BM (*α* = 0). In that case, we take the limit for *α* → 0 represented by the function *u*(*α, t*) in the main text (eq. 5). By substituting *u*(*α, t*) in the expressions for *p_ji_*, *q_ji_* and *r_ji_* (eq. 9) we obtain:

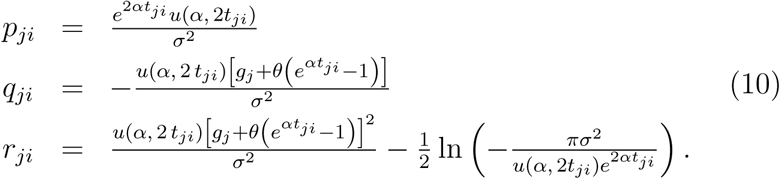 Since *a_i_* < 0 and, for positive *t* and *α* ∈ [0, ∞), *u*(*α, t*) accepts strictly negative values in the interval [-1/*t*, 0), the integral in eq. 9 has a closed form solution:

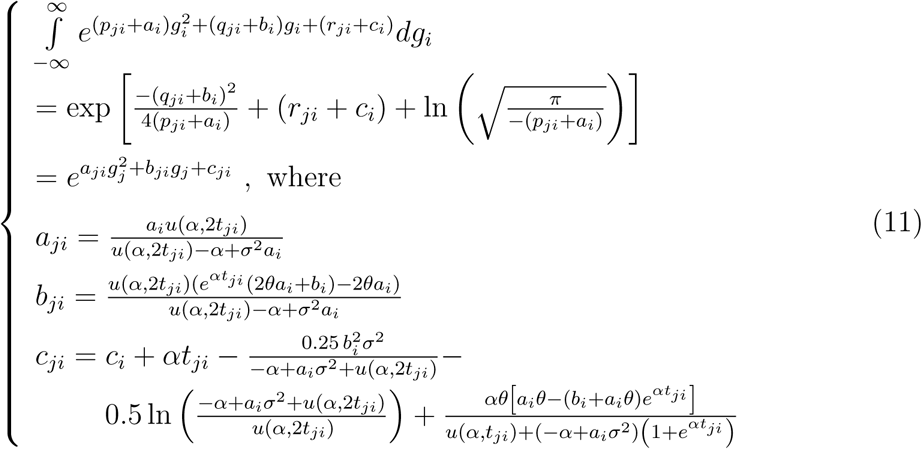 In eq. 11 above, *a_ji_ <* 0 because it is a fraction with a positive nominator and a negative denominator (note that *a_i_ <* 0 by the inductive hypothesis and *u*(*α*, 2*t_ji_*) *<* 0 by definition). Since the vectors **z**_*i*_, *i* ∈ *Desc*(*j*), are conditionally independent given Θ, the conditional pdf of **z***_j_* factorizes as:

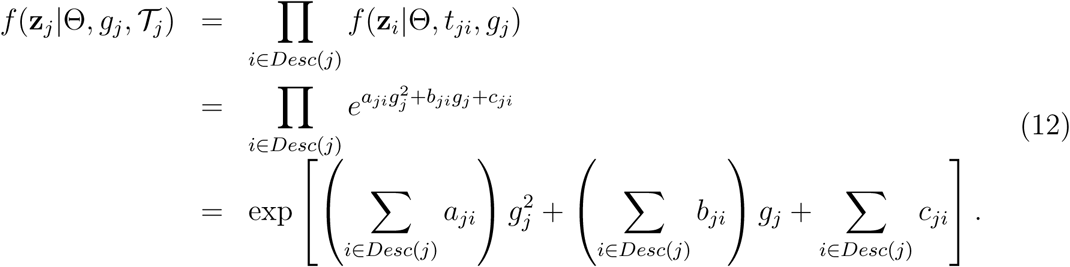 By denoting 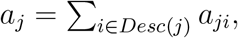, 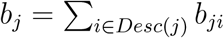 and 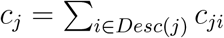 and noticing that *a_j_ <* 0 as a sum of negative terms, we have proven the inductive step and, thus, the theorem.

It can be shown that current pruning implementations (FitzJohn 2012) rely on equivalent formulations of the above theorem. The parallel pruning algorithm differs from these implementations in the ordering of algebraic operations so that they can be performed in parallel for groups of tips or internal nodes rather than consecutively for individual nodes in order of depth-first traversal. This parallelization scheme can also be applied to the generalizaed 3-point representation of the likelihood described in Ho and Ańe (2014), allowing to parallelize the likelihood calculation for some non-Gaussian models such as phylogenetic logistic regression and phylogenetic Poisson regression.

#### Algorithm 1

Parallel pruning

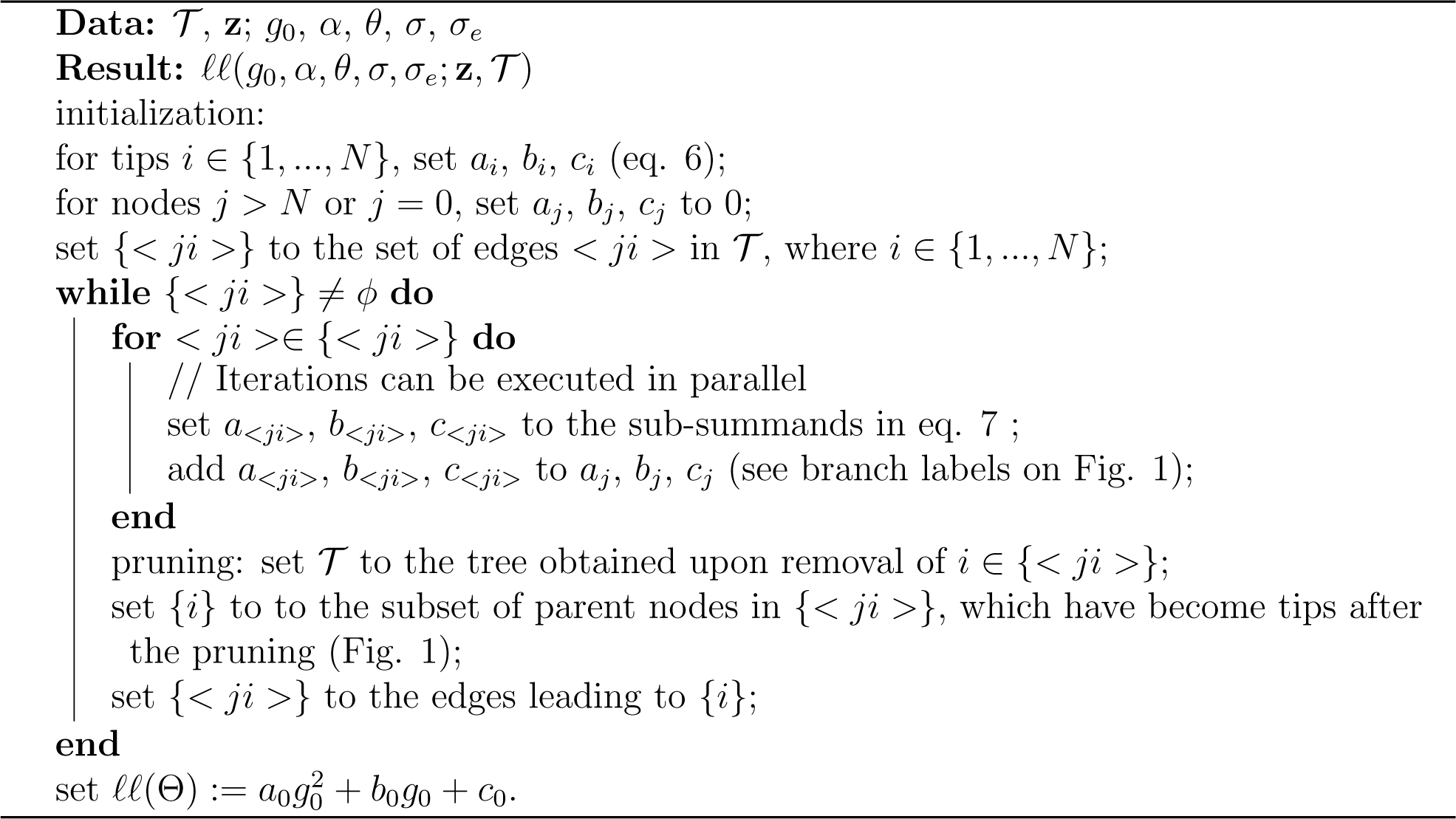

### Applications of the POUMM model

From a modeling perspective, the POUMM can be regarded as a combination of two of its widely used nested models:

- The non-mixed phylogenetic Ornstein-Uhlenbeck (POU) model, (also referred to as the ‘Hansen model’ (Butler and King 2004)), which has been used to infer local phenotypic optima in different phylogenetic clades or across discrete categories of phylogeneticall related species (Butler and King 2004), and to prove the presence of stabilizing selection towards a single stationary peak (SSP) (L. J. Harmon et al. 2010).
- The phylogenetic mixed model (PMM) (Lynch 1991; Housworth, Martins, and Lynch 2004), which has been used to measure phylogenetic heritability (Hodcroft et al. 2014);

Thus, the POUMM combines the applications of the above two models and, as we will show in Results, it is much more accurate and robust in estimating the corresponding parameters. Besides inferring its parameters, the POUMM has several useful properties helping the interpretation of the data and allowing to make predictions about the future trait evolution of the considered population. The properties which we consider represent bijective functions of some of the POUMM parameters. Thus, it is possible to reparametrize the POUMM, so that the model inference is done directly on properties of interest, e.g. the phylogenetic heritability. This is particularly useful for Bayesian inference, since for Bayesian inference priors should be specified for the properties of interest rather than the default POUMM parameters. We call a *parametrization* any numerical bijective function mapping its argument into the default POUMM parameter-space (*< g*_0_*, α, θ, σ, σ_e_ >*).

#### Trait distribution at equilibrium

An interesting property of the POUMM is that, in the limit *t*_0,(_*_ij_*_)_ → *∞*, it defines a stationary normal distribution for the heritable component (*g*) at the tips with mean *θ* and a variance-covariance matrix:

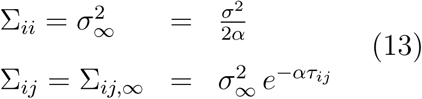

The above property proves useful when there is a prior knowledge that the observed population is at equilibrium, because one can use the trait variance in the population, 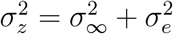 as model parameter. The corresponding parametrization is:

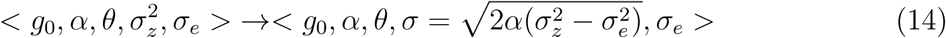

With this parametrization, one can specify an informed prior for 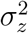 based on empirical estimates on similar data.

Another important aspect of the above property (eq. 13) is that it helps to better understand the selection stregnth parameter *α*. As it turns out, *α* can have two different biological interpretations. Considering the expression for *µ* in table 1, *α* defines the rate of convergence of the population mean towards the long-term optimum *θ*. This rate is bigger for bigger values of *α* and for bigger deviations from *θ*. Thus, *α* is considered as selection strength or rate of adaptation under stabilizing selection. Assuming that the majority of the tips and their mrca’s are far enough from the root, Σ*_ij_* can be viewed as an exponentially decreasing function of the phylogenetic distance *τ_ij_* (eq. 13). Seen from that angle, the parameter *α* can be interpreted as the rate of phenotypic decorrelation between tips, due to genetic drift. When interpreting the results of a model fit, it is important to be aware of this dual interpretation of *α*. In many cases (e.g. in ultrametric macro-evolutionary tree), the only source of information for inferring *α* are the observed differences between the tips in the tree. Thus, in the absence of additional evidence, it can be erronous to assume that the inferred value of *α* informs stabilizing selection and an adaptation rate towards *θ*.

A likelihood ratio test between the ML POUMM and PMM fits can be used to test if the inferred parameter *α* is significantly above 0. As pointed out in the previous paragraph, a significantly positive *α* does not necessarily imply stabilizing selection towards *θ*. Further, it is important to note that the value of *α* can only be interpreted with respect to the time scale of the phylogeny. It can be more intuitive to consider the phylogenetic half-life, 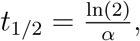 which equals the time it takes for a species entering a new niche to evolve halfway toward its new expected optimum (Hansen 1997).

#### Phylogenetic heritability

The term *phylogenetic heritability*, introduced with the phylogenetic mixed model (PMM) (Housworth, Martins, and Lynch 2004), measures the proportion of phenotypic variance in a population attributable to heritable factors, such as genes, as opposed to non-heritable factors, such as environment and measurement error. Although this concept has been applied mostly in the context of the original PMM, i.e. under the assumption of Brownian motion, the same concept applies to any evolutionary model allowing for the estimation of measurement error (ME) (Hansen and Bartoszek 2012). The *phylogenetic heritability* is defined as the expected proportion of phenotypic variance attributable to *g* at the tips of the tree, 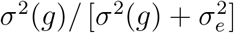 (Housworth, Martins, and Lynch 2004). This definition is a phylogenetic variant of the definition of broad-sense heritability, *H*^2^, from quantitative genetics (Lynch and Walsh 1998). However, in the case of a trait evolving along a phylogeny, the expected genotypic variance, *σ*^2^(*g*), and, therefore, the phylogenetic heritability, are functions of time. Depending on the applicaiton, the following three types of phylogenetic heritability might all be of interest:

- Expectation at the mean root-tip distance 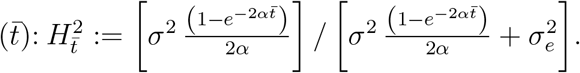. This definition gives rize to three parametrizations where 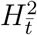 is a free model parameter, while one of the parameters *α*, *σ* or *σ_e_* is a function of 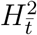 and the two other parameters. These parametrizations can be expressed as:

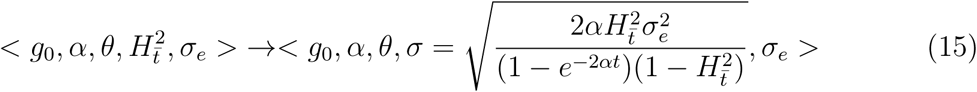

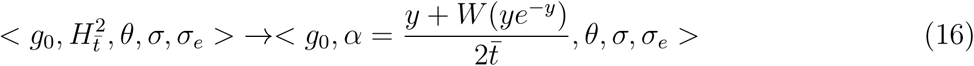

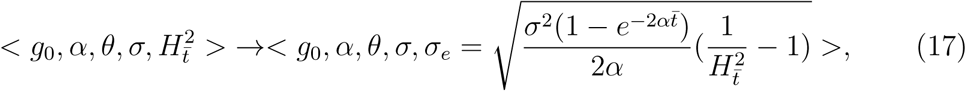

where 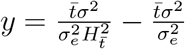 and *W* is the Lambert-W function.
- Expectation at equilibrium of the OU-process (*t* → ∞): 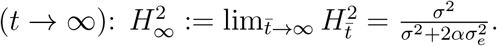. By taking the limit 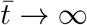 in equations 15, 16 and 17, we obtain the corresponding parametrizations:

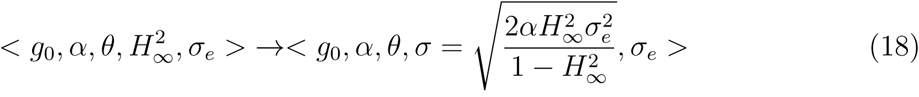

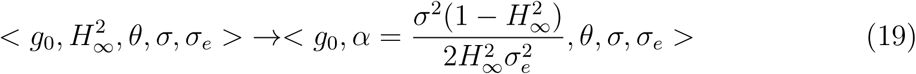

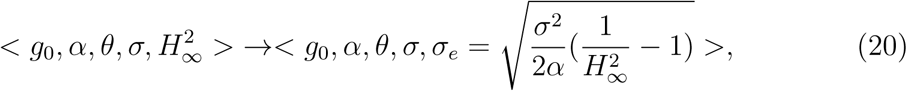
- Empirical (time-independent) version of the heritability based on the sample phenotypic variance 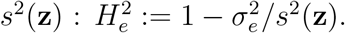. This definition is useful when the tree is non-ultrametric but there is sufficient evidence that the empirical distribution of the trait is stationary along the tree. In this case, *s*^2^(**z**) coincides with the sum of the OU variance at equilibrium and 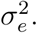 The corresponding parametrization is:

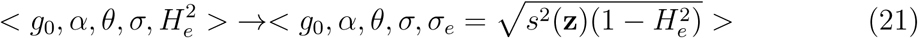

### Implementation

#### Likelihood calculaiton

We tested four implementations of algorithm 1 as follows:

- `R on 1 core`: A serial R-implementation based on operations with numerical vectors in R. This implementation can switch transparently between `double` and `Rmpfr` floating point precision (M. Maechler 2014), thus, guaranteeing numerical stability in cases of extreme parameter values, trait values or branch lengths.
- `C++, Armadillo on 1 core`: A C++ implementation using the library Armadillo (Sanderson and Curtin 2016), through the R-package RcppArmadillo (Eddelbuettel and Sanderson 2014);
- `C++, omp-for on X core(s)`: A serial or parallel C++ implementation where vector elementwise operations are written as C++ for loops and the Open MP preprocessor directive `“#pragma omp for”` is used to parallelize the iterations;
- `C++, omp-for-simd on X core(s)`: A serial or parallel C++ implementation where vector elementwise operations are programmed as C++ for loops and the Open MP preprocessor directive `“#pragma omp for simd”` is used to parallelize and vectorize the iterations.

We compiled the above C++ implementations using version 16.0.0 of the Intel compiler (command icpc with enabled -03 -march = corei7-avx -mavx optons) on

GNU/Linux OS and performed parallelization benchmarks on up to 10 cores on a processor

Intel(R) Xeon(R) CPU E5-2697 v2 @ 2.70 GHz (Results).

#### Possible treatments of g_0_

Recalling that *g*_0_ is an unknown parameter, ℓℓ(Θ) is maximized over *g*_0_ by taking *g*_0_ = − 0.5 *b*_0_*/a*_0_, which is the maximum of eq. 4. During the Bayesian inference, a prior for the parameter has to be specified and it needs to be sampled like all other parameters. Note that, in many cases, e.g. for long ultrametric trees, the likelihood surface can be nearly flat for the parameter *g*_0_, and maximizing over *g*_0_ may result in extremely high or low values.

In these cases, it is better to admit that the data does not inform this parameter, and to exclude it from the free model parameters. This can be done in the following ways:

- assume that *g*_0_ coincides with the long-term optimum *θ*;
- assume a fixed value for *g*_0_;
- Integrate the log-likelihood over *g*_0_ by assuming that it is sampled from a normal distribution such as the OU equilibrium normal distribution with mean *θ* and variance 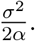 We note, that the latter option may be tricky since the OU equilibrium distribution is not defined for *α* = 0.

#### Fitting the POUMM model

Fitting of the POUMM model was implemented as a pipeline including the following steps, where each step can employ any of the four likelihood implementations mentioned above:

1. Perform three MLE searches using the R-function optim and the L-BFGS-B method (Byrd et al. 1995), starting from three different points in parameter space;
2. Run three MCMC chains as follows: (i) a chain sampling from the prior distribution; (ii) a chain sampling from the posterior distribution and started from the MLE found in step 1; (iii) a chain sampling from the posterior distribution and started from a random point in parameters space.
3. If the parameter tuple of highest likelihood sampled in the MCMC has a likelihood higher than the MLE found in step 1, repeat the MLE search starting from that parameter tuple;

To reduce the number of iterations for MCMC convergence, we use adaptive Metropolis sampling with coerced acceptance rate (Vihola 2012; Scheidegger 2012). By using the R-package foreach (Analytics and Weston 2015), our implementation supports running the MCMC chains in parallel. By comparing the posterior samples from two MCMCs initiated from different starting points, it can be assessed whether the MCMCs have converged to the true posterior. We do this quantitatively by the use of the Gelman-Rubin convergence diagnostic (Brooks and Gelman 1998) implemented in the R-package coda (Plummer et al. 2006). Values of the Gelman-Rubin (G.R.) statistic significantly different from 1 indicate that at least one of the two posterior samples deviates significantly from the true posterior distribution. By visual comparison of posterior density with prior desnity plots, it is possible to assess whether the data contains information differring from the prior for a given sampled parameter.

#### An R-package

We provide the model implementation in the form of an R-package called POUMM. Before model fitting, the user can choose from different POUMM parametrizations and prior settings (function `specifyPOUMM`). A set of standard generic functions, such as `plot, summary, logLik, coef`, etc., provide means to assess the quality of a fit (i.e. MCMC convergence, consistence between ML and MCMC fits) as well as various inferred properties, such as high posterior density (HPD) intervals.

In addition, the POUMM package uses several third-party R-packages: ape (E Paradis, Claude, and Strimmer 2004), data.table (Dowle et al. 2014), coda (Plummer et al. 2006), foreach (Analytics and Weston 2015), ggplot2 (Wickham 2009), GGally (Schloerke et al. 2016), gsl (Hankin 2006) and Matrix (Bates and Maechler 2017).

## Results

### Simulations

To validate the correctness of the Bayesian POUMM implmentation, we used the method of posterior quantiles (S. R. Cook, Gelman, and Rubin 2006). In this method, the idea is to generate samples from the posterior quantile distributions of selected model parameters (or functions thereof) by means of numerous “replications” of simulation followed by Bayesian parameter inference. In each replication, “true” values of the model parameters are drawn from a fixed prior distribution and trait-data is simulated under the model specified by these parameter values. We perform these simulations on a fixed tree of size *N* = 4000. Then, the to-be-tested software is used to produce a posterior distribution of parameters based on the simulated trait-data. Next, the posterior quantiles of the “true” parameter values (or functions thereof) are calculated from the corresponding posterior samples generated by the to-be-tested software. By running multiple independent replications on a fixed prior, it is possible to generate large samples from the posterior quantile distributions of the individual model parameters, as well as any derived quantities. Assuming correctness of the simulations, any statistically significant deviation from uniformity of these posterior quantile samples indicates an error in the to-be-tested software (S. R. Cook, Gelman, and Rubin 2006).

We compared the accuracy of the POUMM to its two nested models - the PMM which restricts *α* = 0 and infers *< g*_0_, *σ*, *σ_e_>*, and the non-mixed Ornstein-Uhlenbeck model (abbreviated as POU) which restricts *σ_e_* = 0 and infers *< g*_0_, *α*, *θ*, *σ>*. Two phylogenetic trees were used for the simulations:

- Ultrametric (BD, *N* = 4000) - an ultrametric birth-death tree of 4000 tips generated using the TreeSim R-package (Stadler et al. 2013, Boskova, Bonhoeffer, and Stadler (2014)) (function call: `sim.bd.taxa(4000, lambda = 2, mu = 1, frac = 1, complete = FALSE)`);
- Non-ultrametric (BD, *N* = 4000) - a non-ultrametric birth-death tree of 4000 tips generated using the TreeSim R-package (Stadler et al. 2013, Boskova, Bonhoeffer, and Stadler (2014)) (function call: `sim.bdsky.stt(4000, lambdasky = 2, deathsky = 1, timesky = 0)`).

Simulation scenarios of 2000 replications were run using the prior distribution *< g*_0_*, α, θ, σ, σ_e_ >∼ N* (5, 25) *×* Exp(0.1) *× U* (2, 8) *×* Exp(0.4) *×* Exp(1). The goal of using this prior was to explore a large enough subspace of the POUMM parameter space, while keeping MCMC convergence and mixing within reasonable time (runtime up to 30 minutes for two MCMCs of 10^6^ adaptive Metropolis iterations at target acceptance rate of 1%). From the above prior, we drew a sample of *n* = 2000 parameter tuples, {Θ^(1)^, …, Θ^(*n*)^}, which were used as replication seeds in two simulation-modes:

- Simulate POUMM - for a given Θ^(*i*)^, simulate genotypic values **g**^(*i*)^(*𝒯*, Θ^(*i*)^) according to an OU-branching process with initial value 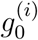 and parameters *α*^(*i*)^, *θ*^(*i*)^, *σ*^(*i*)^. Then add random white noise 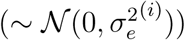 to the genotypic values at the tips, to obtain the final trait values **z**^(*i*)^.
- Simulate PMM - for a given Θ^(*i*)^, reset *α*^(*i*)^to 0, so that the genotypic values come from a BM process, then, repeat the procedure as in mode “Simulate POUMM”. The purpose of implementing this mode was (i) to validate the technical correctness of our PMM implementation by testing for uniformity its posterior quantile distributions; (ii) to obtain an impression of the robustness of the POUMM method to a prior favoring OU (*α >* 0) in the case of true BM processes (*α* = 0).

Combining the two phylogenies with the two simulation modes we obtained four test-scenarios with a total of 4 *×* 2000 = 8000 replications. The resulting posterior quantile distributions for the PMM and POUMM Bayesian fits in each of these scenarios are shown on Fig. 2 for the non-ultrametric and Fig. S1 for the ultrametric tree.

**Figure 2:**
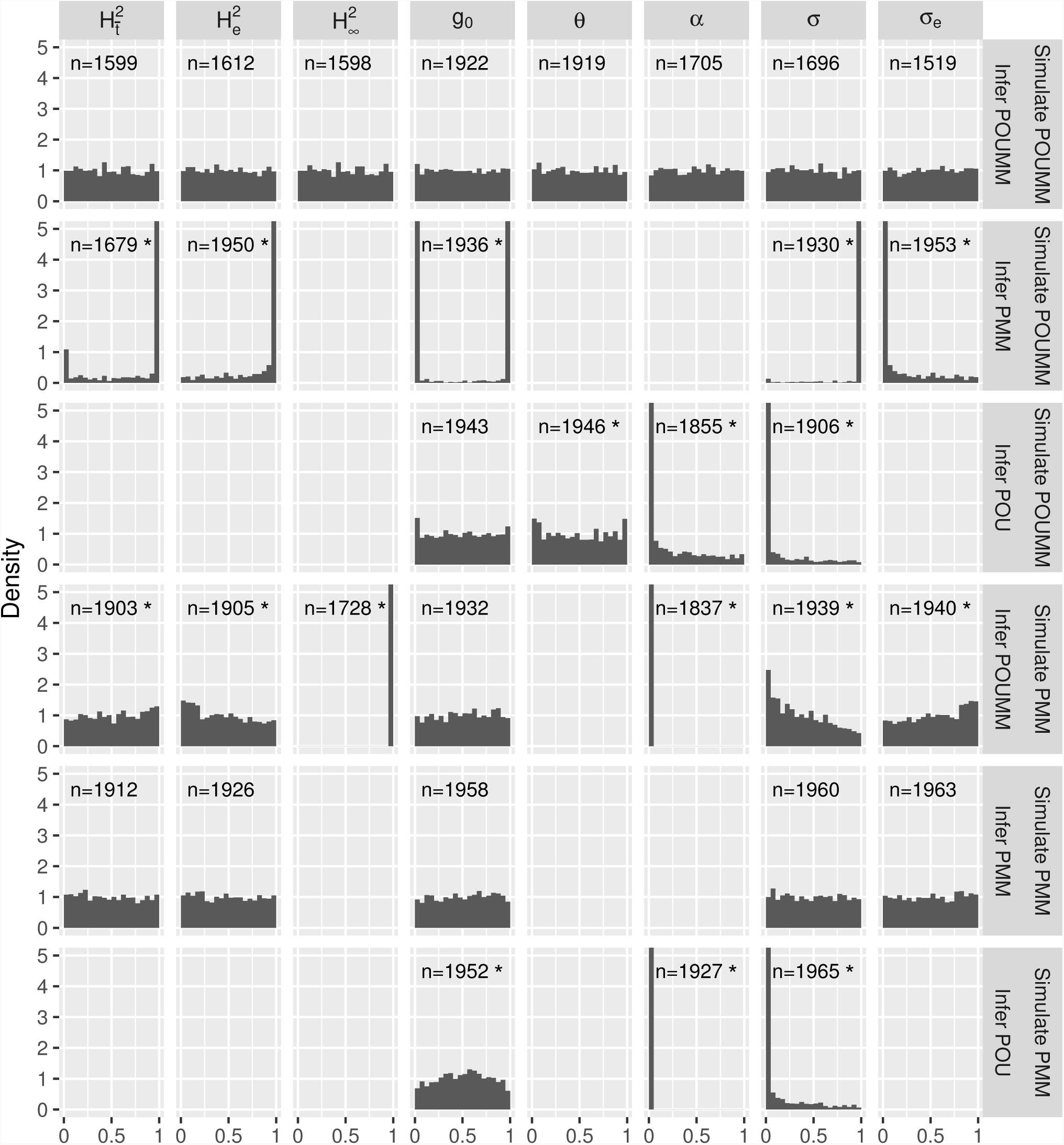
Posterior quantiles from simulation scenarios on a non-ultrametric tree (*N* = 4000). Values tending to 1 indicate that the true value dominates the inferred posterior sample for most of the replications. This means that the model fit tends to underestimate the true parameter. Conversely, values tending to 0 indicate overestimation. The number *n* at the top of each histogram denotes the number of replications out of 2000 which reached acceptable MCMC convergence and mixing after 10^6^ iterations. An asterisk indicates significant uniformity violation (Kolmogorov-Smirnov P-value *<* 0.01). For the analogous results on an ultrametric tree, see Fig. S1.

#### Technical correctness

Both, the PMM and POUMM implementation, generate uniformly distributed posterior quantiles for all relevant parameters when the Bayesian inference has been done on data simulated under the correct simulation mode, i.e. “Simulate PMM” for PMM and “Simulate POUMM” for POUMM. This is confirmed visually by the corresponding histograms (Fig. 2 and Fig. S1), as well as statistically, by a non-significant p-value from a Kolmogorov-Smirnov uniformity test at the 0.01 level. This observation validates the technical correctness of the software.

#### Robustness to model misspecification

Robustness results discussed in this section are all visualized in Fig. 2. When fitting POUMM to data simulated under PMM (*α* = 0), there is a tendency to infer a positive *α*, overestimating *σ* and underestimating *σ_e_*. The cause for this is the wrong prior for *α* pulling it away from 0. Thus, we recommend to always test the hypothesis *α* = 0, e.g. through a likelihood ratio test between the maximum likelihood PMM and POUMM fits. The deviation from uniformity is far less pronounced for 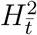 (posterior quantiles tending slightly to 1) and 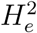 (posterior quantiles tending slightly to 0).

When fitting PMM to simulations under POUMM, there is a highly significant deviation from uniformity of the posterior quantiles for all parameters and derived _quantities. The fact that most posterior quantiles for 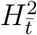 and 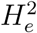 are at the extremes of the_ histogram is indicative for a systematic negative or positive bias in the inferred parameters. These results indicate that the PMM can be a very unstable erronous estimator of phylogenetic heritability when the data violates the Brownian motion assumption.

When fitting POU (*σ_e_* = 0) to simulations including environmental contribution (both simulation modes), there is a strong tendency to overestimate *α* and *σ*. This indicates that POU-inference ignoring non-heritable contributions is prone to false positive tests for stabilizing selection. The next section provides empirical evidence for this prediction.

### Analysis of body weight evolution in mammals

We analysed phylogenetic and body weight data from 227 Rodentia, 138 Chiroptera and 122 Soricomorpha species. An ultrametric tree composed of three monophyletic groups of the above mammal orders was extracted from a previously published supertree of 4510 extant mammal species (Bininda-Emonds et al. 2007) and the body weights have been taken from (Raia, Carotenuto, and Meiri 2010). The orders Rodentia, Chiroptera and Soricomorpha represented the largest groups of species in the supertree with available body weight measures. To extract the phylogeny from the supertree, we used the function “‘drop.tip“‘ from the R-package “‘ape“‘ (E Paradis, Claude, and Strimmer 2004), removing all species of other orders and/or species without available body weight (fig. 3). Further, because all of the phylogenetic models discussed here assume that the trait values at the tips of an ultrametric tree are samples from a normal distribution, we dropped 10 Rodentia, 15 Chiroptera, 4 Soricomorpha species, which appeared as “giants” compared to other species in their corresponding orders and visibly distorted the normality of the data (fig. 3, Discussion). Upon the above filtering, we obtained four trees as follows:

- Rodentia: *N* = 217, 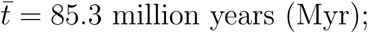
- Chiroptera: *N* = 123, 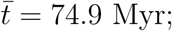
- Soricomorpha: *N* = 118, 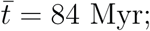
- All (combining the three above trees): *N* = 458, 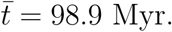

**Figure 3:**
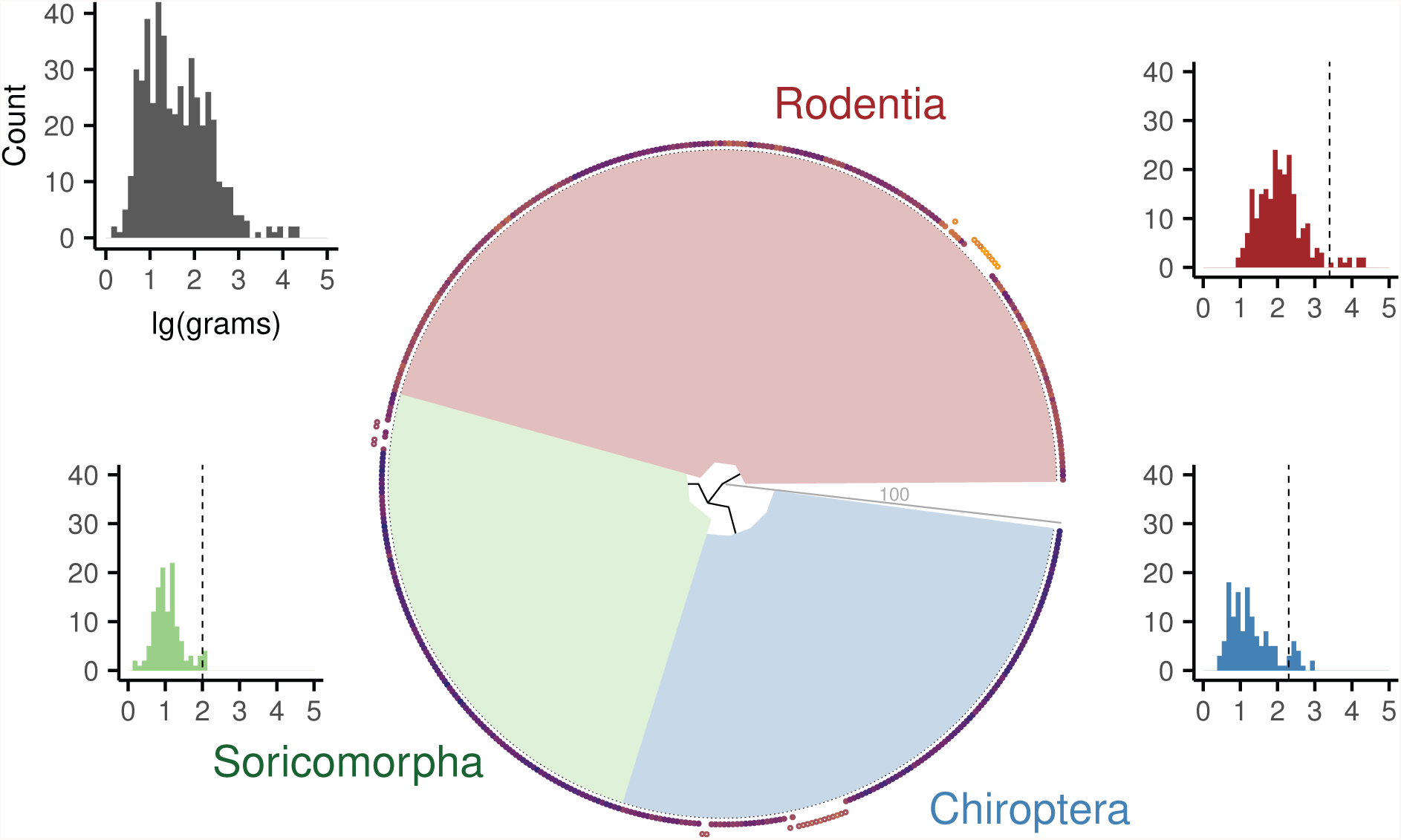
Phylogenetic tree and body weights of 227 Rodentia, 138 Chiroptera and 122 Soricomorpha species (extracted from mammal supertree published in Bininda-Emonds et al. (2007)). Colored bullets represent color-coded lg(body-mass) (blue: low, orange: high, data from Raia, Carotenuto, and Meiri (2010)); outliers (“giants”) species are shown as colored circles positionend slightly outwards; body weight distributions are represented as colored histograms (corresponding to each order), a grey histogram in the top-left corner representing the total body weight distribution across the three orders, including outliers. Dashed vertical bars represent the outlier cut-off for each order. A grey line from the root to the tips indicates the time-scale (length) of the tree in million years (Myr).

For these four trees, we compared the maximum likelihood fits of the following five models:

- PBM / brown: Brownian motion assuming full phylogenetic heritability (*σ_e_* = 0). For this model, we used our implementation based on the POUMM package (parametrization < *θ*, *σ* > → < *g*_0_ ≡ *θ*, *α* = 0, *θ*, *σ*, *σ_e_* = 0 >) and the implementation in the R-package ouch (function brown). The MLEs for the two implementations were matching exactly, except for the tree “All”, where the brown function returned maximum log-likelihood of -∞.
- POU(*∫ g*_0_) / hansen: Ornstein-Uhlenbeck assuming full phylogenetic heritability (*σ_e_* = 0). For this model, we tested our implementation based on the POUMM package (parametrization 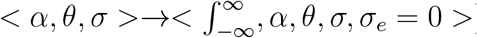) and the implementation in the R-package ouch v2.9.2, function hansen with one global selection regime (Butler and King 2004). By the notation 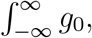 we mean that the OU-likelihood is defined as the expectation of the conditional likelihood on *g*_0_, assuming that the distribution of *g*_0_ is the long-term equilibrium (stationary) OU-distribution. The log-likelihood values and the MLEs for the two implementations were matching up to the 5th decimal digit (table 2).
- POU: Ornstein-Uhlenbeck assuming full phylogenetic heritability (*σ_e_* = 0) and substituting the value of *θ* for the parameter *g*_0_. While the only difference of this model with the Hansen model above is that it replaces integration over *g*_0_ with a concrete value, this model includes PBM/brown as a special case (*α* = 0) and, therefore, one can use a likelihood-ratio test for model selection. For this model, we tested our implementation based on the POUMM package (parametrization < *θ*, *σ* > → < *g*_0_ ≡ *θ*, *α* = 0, *θ*, *σ*, *σ_e_* = 0 >). The resulting maximum log-likelihoods and MLEs were very close to the hansen estimates (table 2).
- PMM: The phylogenetic mixed model (Lynch 1991; Housworth, Martins, and Lynch 2004), i.e. Brownian motion plus error (*σ_e_* ≥ 0). For this model, we used our implementation based on the POUMM package with parametrization 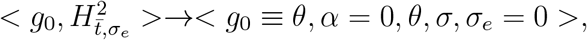 where the parameter *σ* is calculated from eq. 15, after setting *α* to 0.
- POUMM: The phylogenetic Ornstein-Uhlenbeck mixed model. For this model, we used our implementation based on the POUMM package with parametrization 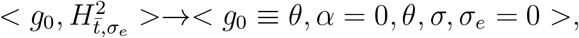 where the parameter *σ* is calculated from eq. 15.

**Table 2:** Model selection criteria and MLEs for five phylogenetic models fitted to mammal body-weight data. The abbreviation LR refers to a likelihood ratio test statistics against a nested null-model (PBM or PMM); asterisks denoting significant p-values: * < 0.05, ** < 0.01, * * * < 0.001. The best (the lowest) AICc values for each tree are shown in bold. Fixed parameters are written in grey. The parameters *g*_0_ and *σ_e_* are not shown, since *g*_0_ is fixed or integrated over (see text) and the positive value of *σ_e_* can be calculated from the phylogenetic heritability 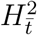. The half-life *t*_1/2_ is given in Millions of years (Myr).

| Model | $N$ | dof | log-lik. | LR vs PBM | LR vs PMM | AICc | MLE:<br>$< \alpha \times 100, \theta, \sigma \times 10, H_t^2, t_{1/2} >$ |
| --- | --- | --- | --- | --- | --- | --- | --- |
| <u>Rodentia</u> |  |  |  |  |  |  |  |
| PBM / brown | 217 | 2 | -66.33 | 0 | - | 136.72 | $< 0, 2.04, 0.81, 1, \infty >$ |
| POU($\int g_0$)/hansen | 217 | 3 | -64.61 | - | - | 135.32 | $< 1.37, 2.03, 0.91, 1, 50.75 >$ |
| POU | 217 | 3 | -64.18 | 4.3* | - | 134.48 | $< 1.23, 2.03, 0.9, 1, 56.46 >$ |
| PMM | 217 | 3 | -63.68 | 5.3* | 0 | **133.48** | $< 0, 2.04, 0.71, 0.96, \infty >$ |
| POUMM | 217 | 4 | -63.46 | 5.75 | 0.45 | 135.11 | $< 0.57, 2.04, 0.77, 0.97, 121.19 >$ |
| <u>Chiroptera</u> |  |  |  |  |  |  |  |
| PBM/brown | 123 | 2 | 2.13 | 0 | - | -0.16 | $< 0, 1.15, 0.59, 1, \infty >$ |
| POU($\int g_0$)/hansen | 123 | 3 | 5.11 | - | - | -4.02 | $< 1.8, 1.12, 0.68, 1, 38.58 >$ |
| POU | 123 | 3 | 5.5 | 6.74** | - | -4.79 | $< 1.67, 1.13, 0.67, 1, 41.61 >$ |
| PMM | 123 | 3 | 11.89 | 19.53*** | 0 | **-17.59** | $< 0, 1.14, 0.43, 0.93, \infty >$ |
| POUMM | 123 | 4 | 11.89 | 19.53*** | 0 | -15.45 | $< 0, 1.14, 0.43, 0.93, \infty >$ |
| <u>Soricomorpha</u> |  |  |  |  |  |  |  |
| PBM/brown | 118 | 2 | -24.24 | 0 | - | 52.58 | $< 0, 1.2, 0.74, 1, \infty >$ |
| POU($\int g_0$)/hansen | 118 | 3 | -23.24 | - | - | 52.68 | $< 1.72, 1.2, 0.84, 1, 40.3 >$ |
| POU | 118 | 3 | -23.05 | 2.37 | - | 52.32 | $< 1.54, 1.2, 0.83, 1, 44.97 >$ |
| PMM | 118 | 3 | -22.19 | 4.09* | 0 | **50.6** | $< 0, 1.2, 0.46, 0.8, \infty >$ |
| POUMM | 118 | 4 | -22.19 | 4.09 | 0 | 52.74 | $< 0, 1.2, 0.46, 0.8, \infty >$ |
| <u>All</u> |  |  |  |  |  |  |  |
| PBM/brown | 458 | 2 | -98.36 | 0 | - | 200.74 | $< 0, 1.56, 0.74, 1, \infty >$ |
| POU($\int g_0$)/hansen | 458 | 3 | -96.42 | - | - | 198.9 | $< 0.8, 1.55, 0.79, 1, 86.78 >$ |
| POU | 458 | 3 | -95.65 | 5.41* | - | 197.35 | $< 0.74, 1.55, 0.79, 1, 94.25 >$ |
| PMM | 458 | 3 | -90.41 | 15.89*** | 0 | **186.88** | $< 0, 1.56, 0.63, 0.96, \infty >$ |
| POUMM | 458 | 4 | -90.4 | 15.9*** | 0.01 | 188.9 | $< 0.07, 1.53, 0.64, 0.96, 1041.29 >$ |

The results from fitting the above models to the four mammal trees are written in table 2. According to best (lowest) Akaike information criterion corrected for finite sample size (AICc), all four trees are unanimous about the best fit being the PMM model. The POUMM method MLE is very similar to the PMM MLE in all cases, but is the second-best model in terms of AICc, because it gets an extra penalty for the extra-parameter *α*, which’s MLE is approximately 0 in all trees. This reveals stronger support for neutral drift evolution (i.e. BM) compared to stabilizing selection acting at the *class* and *order* levels. However, if the model would not account for a non-heritable component in the trait and would assume 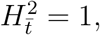 i.e. the PBM/brown, Hansen and POU models, the model selection would turn in favour of the OU-models. This suggests that the conclusion on whether or not stabilizing selection acts on the *class* and *order* levels depends strongly on the inclusion of a non-heritable component in the model, even if this component explains only a small relative proportion of the total phenotypic variance (PMM estimates of phylogenetic heritablity above 93% for all trees, except Soricomorpha).

### Performance

#### Likelihood calculation time

We measured the time for one likelihood calculation on random ultrametric and non-ultrametric trees of up to 100,000 tips running on an Intel(R) Xeon(R) CPU E5-2697 v2 at 2.70 GHz processor with *X* ∈ {1, 2, 4, 6, 8, 10} parallel cores. We compared the serial and parallel pruning implementatoins of the POUMM package to several serial pruning implementations provided in the R-packages geiger (Pennell et al. 2014), and diversitree (FitzJohn 2012). To measure the likelihood calculation times we used the R-package microbenchmark (Mersmann 2015) with argument `times` set to 100. All of the above implementations were compiled from source-code using the R-command `install.packages(‘package-directory’, repos = NULL, type=’source’`), and the same C++ compiler and compiler arguments (version 16.0.0 of the Intel compiler, command `icpc` with options `-03 -march = corei7-avx -mavx`). The resulting average times in milliseconds are shown on fig. 4. On small trees of 100 tips, the fastest implementation was the serial implementation from the package diversitree (0.05 ms) followed by the POUMM `omp-for-simd-on-1-core` (0.07 ms). On trees of 1000 tips, the fastest implementation was the POUMM `omp-for-simd-on-1-core` (0.12 ms) followed by the parallel POUMM implementation `omp-for-simd-on-X-cores` and `omp-for-on-X-cores` (below 0.2 ms). With *N* > 1000 tips, the simd and multicore parallelization resulted in up to 13 times speed-up compared to non-simd serial POUMM C++ implementations (Fig. 5) and speed-ups of up to two orders of magnitude when comparing to serial pruning implementations (Fig. 4 and Fig. 5). These results show that the parallel efficiency tends to increase with *N*, so that on big trees, or in cases of smaller trees but numerous traits, a parallel pruning implementation could potentially achieve 100% parallel efficiency. Also noteworthy is that, thanks to the single instruction multiple data (simd) techology, parallelization is also possible on a single core. This is why, the time for the omp-for-simd implementation on a single core (5 ms for *N* = 100, 000) is several times shorter than the time for the omp-for and the Armadillo based implementations. This also explains why the parallel speed-up is bigger than the number of cores when comparing the `omp-for-simd on X cores` to the non-simd (`omp-for on 1 core`) implementations.

**Figure 4:**
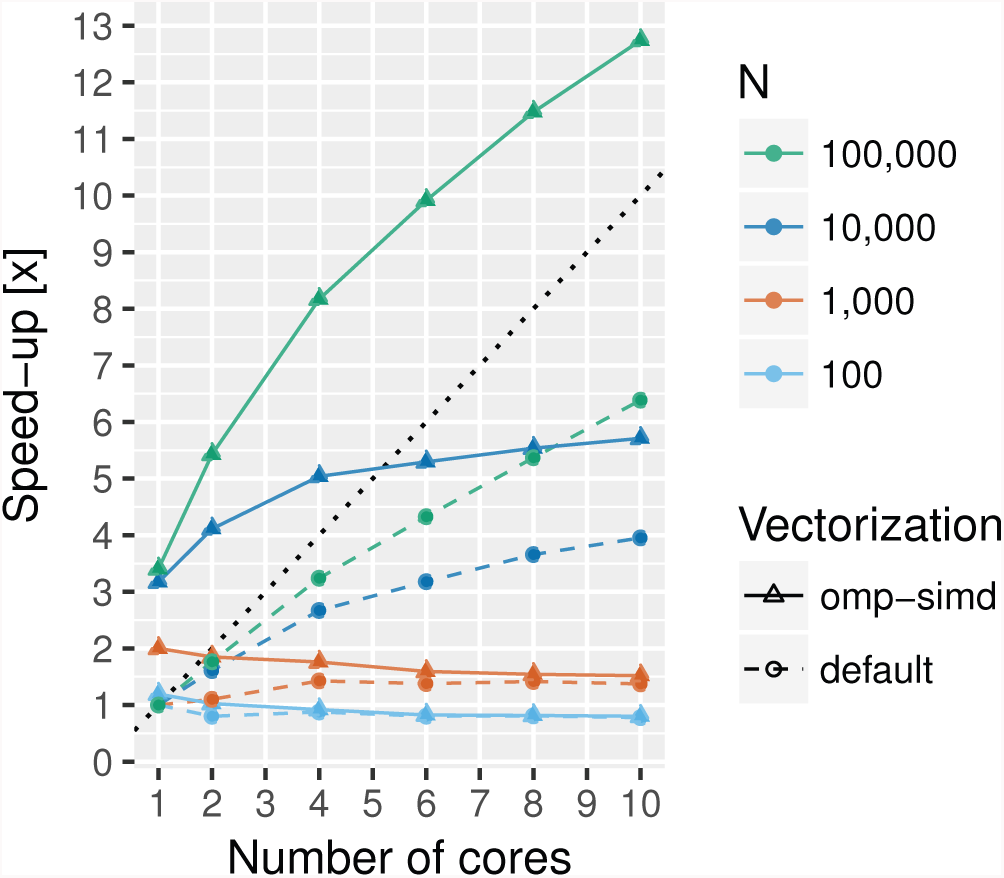
Likelihood calculation times for R and C++ implementations of the pruning algorithm.

**Figure 5:**
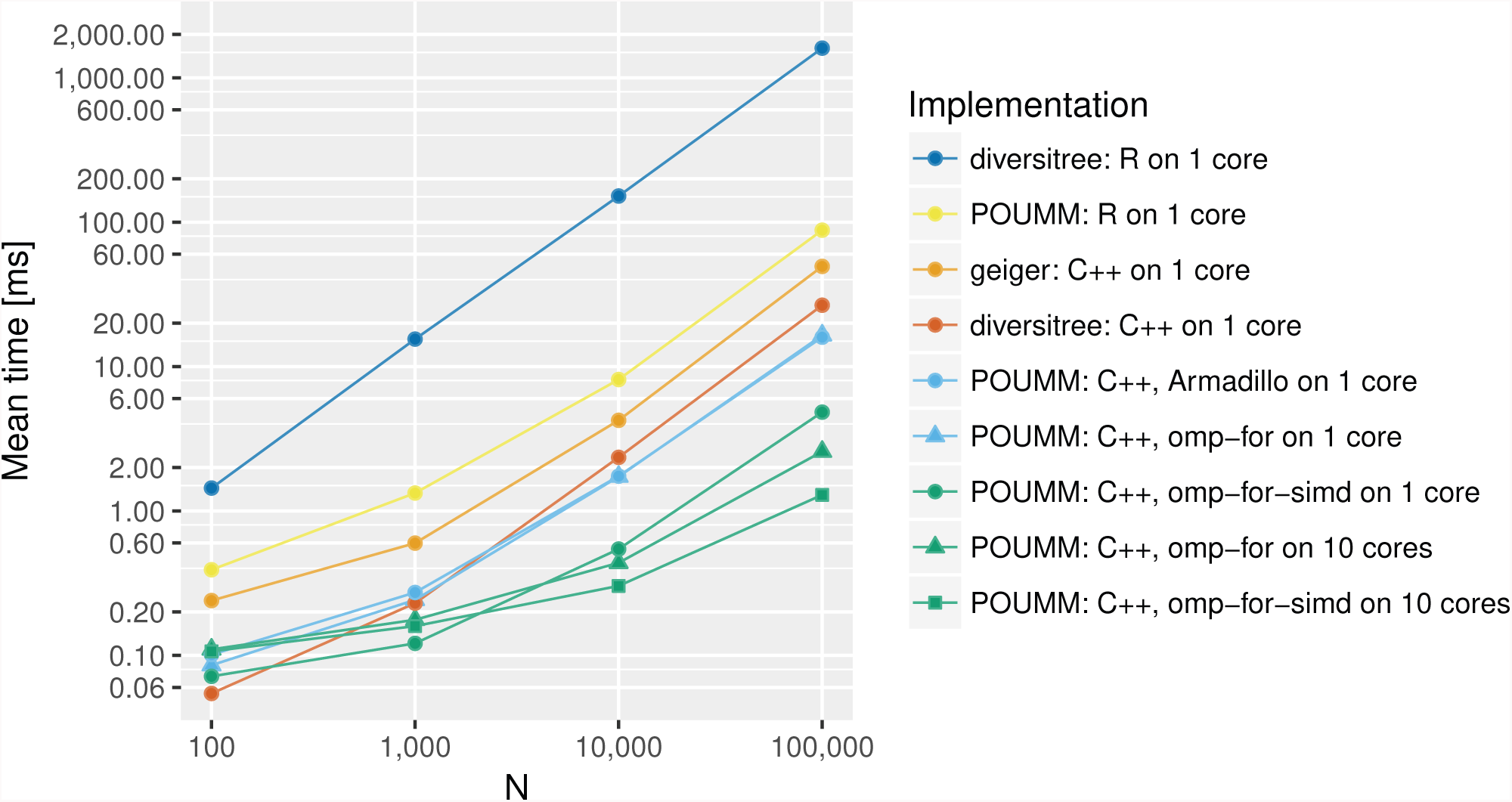
Parallel pruning speed-up on multiple CPU cores. Dashed lines represent the measured parallel speed-up of omp-for implementations with respect to the serial non-simd (omp-for) implementation running on one core; continuous lines represent the parallel speed-up of omp-for-simd implementation with respect to the serial non-simd (omp-for) implementation.

#### Improved MCMC convergence and MLE inference through adaptive Metropolis sampling

To measure the MCMC convergence speed-up from the adaptive Metropolis sampling, we reran one simulation scenario (2000 replications on a non-ultrametric tree of 4000 tips) with disabled adaptation. As a criterion for convergence, we used the absolute difference from 1 of the Gelman-Rubin convergence diagnostic (Brooks and Gelman 1998) (the closer |*G.R.* – 1| is to 0, the better the convergence). When enabling adaptive Metropolis sampling, more than 1600 (80%) of the 2000 replications had reached |*G.R.* – 1| < 0.01 after a million iterations. Conversely, when disabling adaptive Metropolis sampling, less than 300 (15%) of the replications had reached |*G.R.* – 1| < 0.01 after a million iterations (the 80% quantile of |*G.R.* – 1| was equal to 0.57, indicating very power convergence). We also noticed that 1455 out of 2000 replications (73%) of the POUMM inferences with enabled adaptative Metropolis sampling resulted in an improved MLE after running the MCMC chains, compared to 1045 (50%) when disabling adaptation. These observations show that adaptive Metropolis sampling considerably accelerates the MCMC convergence towards the posterior distribution and can be used to improve the MLE inference when using a weak prior or a prior that does not strongly contradict with the evidence (likelihood).

## Discussion

The OU process has been applied as a model for stabilizing selection in macro-evolutionary studies (LANDE 1976; Felsenstein 1988; Hansen 1997; L. J. Harmon et al. 2010; Raia and Meiri 2011). Most of these studies assume that the whole trait evolves according to an OU process, usually disregarding the presence of a biologically relevant non-heritable component. When modelling species trait evolution, a non-heritable component including ecological contribution, measurement error, model residual error and possible inaccucies in the phylogenetic tree may be well justified and may in fact be important to understand the full evolutionary process. In our empirical example, we have shown that the evolution of body weight in three orders of mammals is best described by the sum of a Brownian motion process and a non-heritable white noice (PMM). Except for the Soricomorpha tree, in which most of the species are in an unresolved polytomy descending from a single ancestor (12.6 Myr), the relative contribution of white noice to the trait variance is estimated between 3% to 7%. Strikingly, it is the inclusion or not of this small contribution in the model that decides the choice between a BM or an OU model of evolution.

Recently, the OU process has been applied in a micro-evolutionary context as a model for the evolution of pathogen traits, such as set-point virus load (spVL) during HIV infection (Mitov and Stadler (2016), Blanquart et al. (2017), Bertels et al. (2017), Bachmann et al. (2017)). When modeling pathogen evolution, the branching points in the tree represent transmission events, and the non-heritable component includes the contribution of the host immune system, the environment and the measurement and phylogenetic error. Thus, for pathogens, it is crucial to incorporate *e* in the model in order to quantify the importance of host-versus pathogen factors in trait formation (Alizon et al. 2010; Shirreff et al. 2013). In contrast to our macro-evolutionary example, based on AICc, the above studies have selected an OU process with added white noice (POUMM) as the best model for the evolution of spVL and CD4.

In agreement with our simulations, the above empirical examples provide evidence that mixed models are more appropriate than PBM and POU for modeling the evolution of continuous traits in epidemiology and macroevolution. Note also that the concept of model mixing and phylogenetic heritability is applicable to any phylogenetic model, beyond BM and OU.

Our R-package joins a growing collection of tools implementing phylogenetic OU inference. Among others, these include the R-packages `ape v4.0` (E Paradis, Claude, and Strimmer 2004), `ouch v2.9-2` (Butler and King 2004), `GLSME v1.0.3` (Hansen and Bartoszek 2012), `diversitree v0.9-9` (FitzJohn 2012), `geiger v2.0.6` (Pennell et al. 2014), `surface v0.4-1` (Ingram and Mahler 2013), `mvMORPH v1.0.8` (Clavel, Escarguel, and Merceron 2015), `bayou v1.0.0` (Uyeda, Eastman, and Harmon 2015), `OUwie v1.50` (Beaulieu and OMeara 2016), `phylolm v2.5` (Ho and Ańe 2014), `RPANDA v1.1` (Manceau, Lambert, and Morlon 2016). It may come as a surprise that most of the above package versions do not implement inference of a non-heritable component, i.e. a parameter *σ_e_*. To our knowledge, of the above mentioned package versions only geiger and GLSME allow *σ_e_* to be estimated from the phylogeny. However geiger does not seem to support non-ultrametric trees (mismatching likelihood values on non-ultrametric trees), and GLSME is much slower than the other packages. Most of the above packages allow the specification of a fixed standard measurement error before the model is fit to the data.

This seems useful in macro-evolutionary studies where the standard measurement error can be estimated from the observed empirical variance within a species and the finite sample size. However, estimating a parameter *σ_e_* from the data is still necessary, because measurement error is only one of many non-heritable factors contributing to the trait.

The idea to infer phylogenetic heritability assuming that *g* follows an OU process along the phylogeny has so far been discouraged mainly for interpretational and practical reasons: (i) in biology, individuals get selected based on their whole trait values, rather than the genotypic component *g*; (ii) small ultrametric macro-evolutionary trees do not contain sufficient signal for a simultaneous inference of the OU-and environmental variance (Housworth, Martins, and Lynch 2004). We argue that modeling an OU process on the whole trait value rather than *g* comes at the cost of additional parameters and reduced statistical power, because it necessitates to account for jumps in the trait value at the branching points as well as the unobserved speciation/transmission events along the tree. Conversely, assuming that the OU process acts directly on *g* is mathematically more convenient, because it allows the inference of a single continuous OU-process along the tree, while adding *e* only at the tips of the tree. The implications of these simplified assumptions must be validated through simulations as done, e.g. in toy model simulations in Mitov and Stadler (2016).

Finally, we have shown the gain in speed performance from parallelizing the likelihood calculation of a univariate Ornstein-Uhlenbeck model. A main advantage of the parallel pruning algorithm with respect to sequential pruning implementations, e.g. diversitree (FitzJohn 2012) and geiger (Pennell et al. 2014), is that most of the algebraic calculations are done on vectors instead of single numbers and can be executed in parallel on contemporary SIMD and multicore systems. Languages such as Matlab and R are optimized for vector operations. This also explains why the serial POUMM pruning implementation in R is nearly as fast as serial pruning implementations written in C++ (Fig. 4). The parallelization scheme described here applies to all phylogenetic models where pruning can be used for likelihood calculation. Previously, it has been shown that this class of models spans over all multivariate Gaussian and some non-Gaussian models (Ho and Ańe 2014). It is noteworthy that the performance benefit from parallelization increases with the complexity of the model (i.e. number of observable variables) and the size of the data (number of observations, *N)*.

Another feature of the parallel pruning alogrithm is that it can be applied when parameters of the model change through time or across clades, e.g. when the values of the parameters are functions of geological time, geographic location or environment. Following this approach, the lineages of the tree can be cut into segments associated with different model regimes. These applications suggests that the parallel pruning algorithm has the potential to meet the challenges of increasing model complexity and volumes of data in comparative phylogenetic analysis.

## Supplementary Material

Supplementary figures are available online. Data from the performance benchmarks, simulations and the analysis of mammal body weight is available on the dryad database. The POUMM package and user guide is available at https://CRAN.R-project.org/package=POUMM.

## Acknowledgements

We thank Dr. Krzysztof Bartoszek for valuable insights on the Ornstein-Uhlenbeck process.

